# SARS-CoV-2 in the Republic of Guinea: Fragment and Whole-Genome Sequencing, Phylogenetic Analysis

**DOI:** 10.1101/2021.07.28.454098

**Authors:** A.A. Kritsky, Ya.M. Krasnov, M. Keita, S. Keita, A.V. Fedorov, A.P. Shevtsova, N.P. Guseva, E.V. Kazorina, E.A. Sosedova, A.D. Katyshev, E.A. Naryshkina, E.V. Kolomoets, S.A. Shcherbakova, A.Yu. Popova, V.V. Kutyrev

## Abstract

Genetic diversity of SARS-CoV-2 isolates circulating in the Republic of Guinea in May and June 2020, as well as in March 2021, has been demonstrated using fragment (S gene) and whole genome sequencing of 14 strains. Analysis of nucleotide sequences and phylogenetic constructs make it possible to divide the studied strains into 3 groups. Comparison of the obtained data with the already available epidemiological data proves the initial importation of COVID-19 from Western European countries, and also demonstrates four independent import routes in two time periods (March 2020 and no later than March 2021).

## Introduction

The pandemic of the new coronavirus infection, COVID-19, the etiological agent of which is the SARS-CoV-2 coronavirus, has affected absolutely all states of the world. As of May 27, 2021, 169,094,726 laboratory-confirmed cases of the disease were registered in the world, 3,512,510 of which were fatal [1]. In African states, 4,830,593 cases were reported, including 129,620 deaths. The Republic of Guinea accounted for 23,080 cases, including 160 deaths.

Having spread around the world, since December 2019, the etiological agent of COVID-19 has undergone various changes in the genome in the course of its evolution, producing many strain genovariants [2, 3, 4]. It is believed that strains with changes in the gene of the spike glycoprotein (“S gene”), especially in the region encoding the receptor binding domain (RBD), are of epidemic significance. [5, 6] Changes in the structure of the spike protein are primarily associated with an increase in the infectiousness of the causative agent of COVID-19, and also contribute to the escape of the virus from the immune response [7, 8, 9].

Among the whole variety of genetic variants of SARS-CoV-2, as of May 31, 2021, the World Health Organization recognized the following variants of the virus as epidemically significant [10], for which an increase in transmissibility, severity of the disease, as well as a decrease in sensitivity to established therapy regimens was proven [11, 12, 13, 14]:

- Alpha – B.1.1.7 (501Y.V1) was first detected in samples collected in the UK in September 2020. The Alpha variant has been registered in 114 countries.
- Beta – B 1.351 (501.V2), first described in South Africa in December, 2020. This strain has become widespread in 92 countries of the world.
- Gamma – P1 (501Y.V3). It was first registered in Brazil in December, 2020. It has spread in 49 states.
- Delta – B.1.617.2 (478K), first discovered in India in October, 2020. Registered in 63 states.

Today, intensive work is being carried out on sequencing and bioinformatic analysis of the genomes of SARS-CoV-2, in order to identify new gene variants as well as to establish pathways for the spread of existing ones, since epidemically significant gene variants pose a serious threat to public health. However, for developing countries, sequencing of the detected strains of SARS-CoV-2 is problematic due to economic reasons, poor logistics, and other factors [15, 16]. In this regard, the Republic of Guinea is no exception. Within the framework of the Russian-Guinean scientific and technical cooperation fragment and whole genome sequencing of SARS-CoV-2 genovariants circulating in the territory of the Republic of Guinea has been performed.

The main task of our study was to investigate the nucleotide sequences of the S gene, as well as the complete genomes of SARS-CoV-2 strains isolated in the Republic of Guinea at different time periods, in order to study their genetic diversity, as well as to establish their origin.

To achieve this goal, the following objectives were identified:

1. Fragment sequencing (S gene).
2. Whole genome sequencing.
3. Construction of phylogenetic trees based on the sequencing of both the S gene and complete genomes.
4. Bioinformatic analysis of the results.

## Materials and methods

Specimens – nasopharyngeal swabs were collected from suspected cases and contact persons as part of the public health emergency response of the Ministry of Health in the Republic of Guinea.

Isolation of RNA from specimens was carried out using a kit for the isolation of nucleic acids “RiboPrep” (“InterLabService”, Russia). Detection of the SARS-CoV-2 genetic material was carried out using the RT-PCR kits: “Detection Kit for 2019 Novel Coronavirus (2019-nCov) RNA (PCR-Fluorescence Probing)” (Da An Gene Co. Ltd., China), “Novel Coronavirus (2019-nCov) RT-PCR Detection Kit” (Fosun Pharma, China), “Novel Coronavirus (2019-nCov) Nucleic Acid Diagnostic Kit (PCR-Fluorescence Probing)” (SanSure Biotech Inc., China) in accordance with the manufacturers’ protocols. The polymerase chain reaction was run on a RotorGene Q (Qiagene, Germany). Reverse transcription of RNA into cDNA was performed using the “Reverta-L kit” (InterLabService, Russia).

Sequencing of complete genomes was carried out using semiconductor sequencing technology on the Ion S5TM XL System platform (Thermo Fisher Scientific, USA) according to the manufacturer’s protocol. The primary data processing was carried out using the Torrent Suite Software version 5.12.1 (Thermo Fisher Scientific, USA). The genome was assembled through mapping the filtered reads to the reference genome (Wuhan-Hu-1 (NC_045512.2)) using the BWA software version 07.17. S-gene sequencing was carried out according to the Sanger method on an AB 3500xl (Applied Biosystems, USA) using Data Collection v3.1 software. Alignment of the nucleotide sequences of both complete genomes and the S gene was carried out using the MAFFT v.7 program (https://mafft.cbrc.jp/alignment/server/). The search for single nucleotide polymorphisms (SNPs) in the compared genomes was performed using the Snippy 2.0 program (https://github.com/tseemann/snippy). Comparative phylogenetic analysis was conducted using the BioNumerics 7.6 software package (Applied Maths NV, Belgium) and the Maximum Parsimony algorithm.

In total, 14 samples obtained in different time periods and from different regions of the Republic of Guinea were analyzed. Whole genome sequencing was carried out for 4 samples, in other 10 samples only the S gene was sequenced. The obtained nucleotide sequences were deposited in the international databases NCBI GenBank. Data on the studied samples are presented in Table 1.

**Table 1.** Specimens included in the study.

| № | Sample ID | Prefecture | Sub-Prefecture | Settlement | Date of sampling | Genotype | Type of sequencing | NCBI GenBank accession |
| --- | --- | --- | --- | --- | --- | --- | --- | --- |
| 1. | C1 | Kindia | Kindia | Kindia | 14.05.2020 | GR | Whole-genome | MZ619091 |
| 2. | C2 | Boke | Kamsar | Kamsar | 13.05.2020 | GR | Whole-genome | MZ622025 |
| 3. | C3 | Boke | Kolaboui | Kolaboui | 14.06.2020 | G | Whole-genome | MZ604340 |
| 4. | C4 | Boke | Sangaredi | Sangaredi | 26.06.2020 | G | Whole-genome | in submission |
| 5. | E10 | Kindia | Kindia | Souguetta | 02.03.2021 | 202012/01, B.1.1.7, Alpha | S gene | MZ604347 |
| 6. | E11 | Kindia | Kindia | Sinania | 10.03.2021 | Alpha alt* | S gene | MZ604348 |
| 7. | E12 | Kindia | Kindia | Damakania | 10.03.2021 | unknown | S gene | MZ604350 |
| 8. | E13 | Kindia | Kindia | Damakania | 10.03.2021 | unknown | S gene | MZ604349 |
| 9. | E14 | Kindia | Kindia | Yabara | 10.03.2021 | Alpha alt* | S gene | MZ604351 |
| 10. | E15 | Kindia | Kindia | Damakania | 10.03.2021 | 202012/01, B.1.1.7, Alpha | S gene | MZ604352 |
| 11. | E16 | Kindia | Kindia | Comoyah Plateau | 12.03.2021 | Alpha alt* | S gene | MZ604353 |
| 12. | E17 | Kindia | Kindia | Ferefou 1 | 16.03.2021 | 202012/01, B.1.1.7, Alpha | S gene | MZ604357 |
| 13. | E18 | Kindia | Kindia | Gadawawa | 17.03.2021 | 202012/01, B.1.1.7, Alpha | S gene | MZ604356 |
| 14. | E20 | Kindia | Kindia | Sambaya | 20.03.2021 | unknown | S gene | MZ604358 |

\*Alpha alt – samples that have all of the Alpha variant marker mutations, but do not have specific deletions

## Results and discussion

The first laboratory-confirmed case of new coronavirus infection in the Republic of Guinea was registered on March 12, 2020 in the capital city, Conakry, in a citizen of the European Union, who arrived from France aboard the airplane shortly before the registration of the disease [17]. In total, in March 2020, until the closure of international traffic (in accordance with the Order of the Government of Guinea, dated March 23, 2020 air traffic was suspended for an indefinite period, and since March 27, the land borders [18] were also closed) 22 cases of the disease, 9 of which were reliably imported, were reported. Among those 9 cases, 8 were related to import from France, and one from Italy [17]. The subsequent spread was associated with community transmission of the virus, up to the opening of the borders in August 2020. It should be noted that, apart from import by aircraft, the likelihood of bringing the virus in by crew members of sea vessels was also not excluded, in view of the presence of large port cities in Guinea – Conakry, Boke and Boffa, the importance of which as a “COVID-19 entrance gates” was emphasized in “National COVID-19 Response Plan” [19].

Throughout the entire period, specialists from the Russian-Guinean Research Center for Epidemiology and Prevention of Infectious Diseases were directly involved in diagnostic and anti-epidemic measures in Guinea.

The number of detected COVID-19 cases between March 2020 and June 2021 was shifting and underwent fluctuations. Peak values were noted in May, mid-June and July, August, October 2020 and March 2021 [20]. We examined samples collected in Kindia and Kamsar in May, 2020 (1 sample each), June 2020 (Kolaboui and Sangaredi cities, 1 sample each), as well as in 8 settlements of Kindia sub-prefecture (10 samples) in March 2021. It is important to note that Kamsar, Kolaboui and Sangaredi settlements are located in Boke Prefecture, a region with a developed mining industry, traditionally home to a large number of employees of foreign mining companies involved in the extraction of minerals. The sub-prefecture of Kindia, from where the rest of the studied samples came, is a major economic center as well as a key transport hub [21]. It is through Kindia that the internal migration of the population from the capital to the inner regions of the country and vice versa, as well as to neighboring states, is carried out. In total, 107 laboratory-confirmed cases of COVID-19 were detected in the Kindia sub-prefecture in March 2021, so we examined 9,3% of isolates from this territory over March.

Bioinformatic analysis showed that the studied samples were not genetically homogeneous. Analysis of the nucleotide sequence of the S gene, carried out for all 14 samples, allows them to be divided into 3 groups. The first includes samples (C1, C2, C3, C4) taken in May-June 2020 in the prefectures of Boke (Kamsar, Kolaboui, Sangaredi) and Kindia (Kindia). All of them have the D614G mutation, and the C3 isolate also has an additional Cys1235Phe mutation, but they do not belong to the known genovariants of epidemic significance. Strains of the same genotype have been described for the states of Western Europe, Asia, and North America. Importantly, sample C1 is one of the first laboratory-confirmed cases of COVID-19 in the Kindia sub-prefecture.

The second group of samples included 7 isolates dated March 2021, with the Alpha genotype (B.1.1.7), or close to it. Four samples (E10, E15, E17, E18) had the entire set of marker mutations and a number of additional mutations. Thus, E10 isolate had an additional mutation Glu619Lys, E15 – Pro9Ser, E17 had a Gln965His, and E18 had Ser12Cys, Thr76Asn, and Gln965His. All of them were registered in different settlements of the Kindia sub-prefecture – Souguetta (E10), Damakania (E15), Ferefou I (E17), Gadawawa (E18). Three more samples (E11, E14, E16), despite the presence of all single substitutions marker for the Alpha genotype, did not have its marker amino acid deletions in the positions 69, 70, 144, while samples E11 and E14 had an additional mutation Asp138Tyr. The samples also have different geographic confinedness within the sub-prefecture of Kindia – Sinania (E11), Yabara (E14), Commoyah Plateau (E16).

The third group included three samples (E12, E13, E20, Kindia sub-prefecture, March 2021) carrying a common unique set of mutations in the S gene: K182R, L452R, T478K, D614G, P681H, D796Y. Interestingly, mutations L452R, T478K are one of the markers for the Delta variant (B.1.617.2); however, other differences do not allow assigning the studied isolates to the Delta genovariant. Samples E12 and E13 were utterly identical to each other and were obtained from settlement Damakania (sub-prefecture of Kindia). Sample E20, which differs from E12 and E13 by 4 SNPs, but is in the same group with them, was isolated in another settlement, Sambaya. Strains of a genotype similar to the third group in the international NCBI GenBank database are represented by only a few specimens from France, Luxembourg and the United Kingdom.

It is important to note that the isolates of the second and third groups, despite their genetic heterogeneity, circulated in the territory of a common sub-prefecture (Kindia) in the same time period (biomaterial samples were taken between March 2 and March 20, 2021), which indicates independent pathways of importation of infection into the territory. The results of S-gene sequencing of isolates of all groups are presented in Table 2.

**Table 2.**
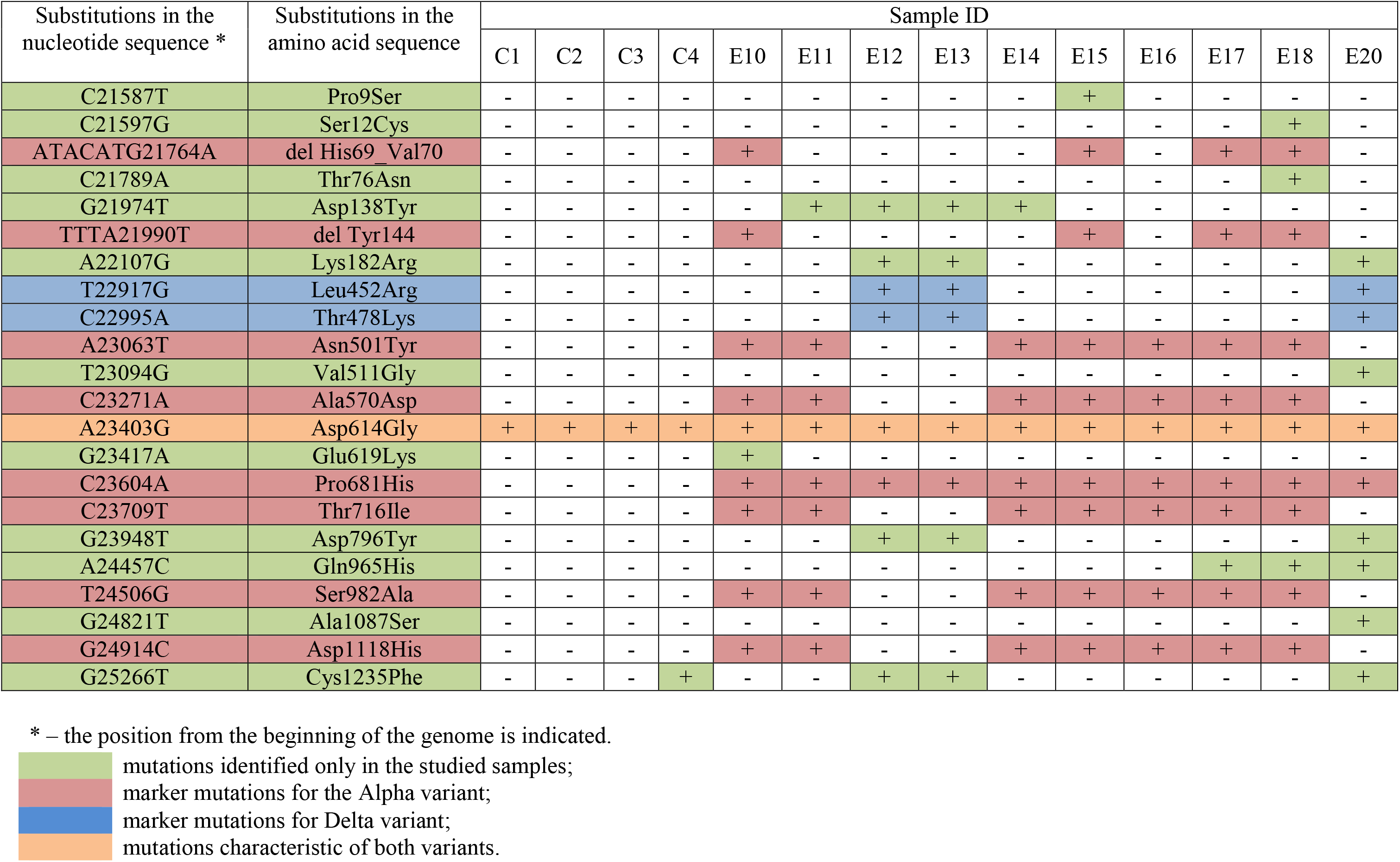
Non-synonymous mutations in the S gene in the studied samples.

| Substitutions in the nucleotide sequence * | Substitutions in the amino acid sequence | Sample ID |  |  |  |  |  |  |  |  |  |  |  |  |  |
| --- | --- | --- | --- | --- | --- | --- | --- | --- | --- | --- | --- | --- | --- | --- | --- |
|  |  | C1 | C2 | C3 | C4 | E10 | E11 | E12 | E13 | E14 | E15 | E16 | E17 | E18 | E20 |
| C21587T | Pro9Ser | - | - | - | - | - | - | - | - | - | + | - | - | - | - |
| C21597G | Ser12Cys | - | - | - | - | - | - | - | - | - | - | - | - | + | - |
| ATACATG21764A | del His69 Val70 | - | - | - | - | + | - | - | - | - | + | - | + | + | - |
| C21789A | Thr76Asn | - | - | - | - | - | - | - | - | - | - | - | - | + | - |
| G21974T | Asp138Tyr | - | - | - | - | - | + | + | + | + | - | - | - | - | - |
| TTTA21990T | del Tyr144 | - | - | - | - | + | - | - | - | - | + | - | + | + | - |
| A22107G | Lys182Arg | - | - | - | - | - | - | + | + | - | - | - | - | - | + |
| T22917G | Leu452Arg | - | - | - | - | - | - | + | + | - | - | - | - | - | + |
| C22995A | Thr478Lys | - | - | - | - | - | - | + | + | - | - | - | - | - | + |
| A23063T | Asn501Tyr | - | - | - | - | + | + | - | - | + | + | + | + | + | - |
| T23094G | Val511Gly | - | - | - | - | - | - | - | - | - | - | - | - | - | + |
| C23271A | Ala570Asp | - | - | - | - | + | + | - | - | + | + | + | + | + | - |
| A23403G | Asp614Gly | + | + | + | + | + | + | + | + | + | + | + | + | + | + |
| G23417A | Glu619Lys | - | - | - | - | + | - | - | - | - | - | - | - | - | - |
| C23604A | Pro681His | - | - | - | - | + | + | + | + | + | + | + | + | + | + |
| C23709T | Thr716Ile | - | - | - | - | + | + | - | - | + | + | + | + | + | - |
| G23948T | Asp796Tyr | - | - | - | - | - | - | + | + | - | - | - | - | - | + |
| A24457C | Gln965His | - | - | - | - | - | - | - | - | - | - | - | + | + | + |
| T24506G | Ser982Ala | - | - | - | - | + | + | - | - | + | + | + | + | + | - |
| G24821T | Ala1087Ser | - | - | - | - | - | - | - | - | - | - | - | - | - | + |
| G24914C | Asp1118His | - | - | - | - | + | + | - | - | + | + | + | + | + | - |
| G25266T | Cys1235Phe | - | - | - | + | - | - | + | + | - | - | - | - | - | + |

\* – the position from the beginning of the genome is indicated.
mutations identified only in the studied samples; marker mutations for the Alpha variant; marker mutations for Delta variant; mutations characteristic of both variants.

Phylogenetic analysis using the nucleotide sequence of the S gene showed that the samples of the first group (obtained in 2020) were greatly distanced from the isolates of 2021, and were in the same clade with the isolates circulating in 2020 in many countries of the world, without a clearly defined territorial affiliation, which confirms the data on the importation of COVID-19 into Guinea from abroad. The genovariants of the second group also formed a separate clade, being located together with the typical variants of the Alpha line. Samples of the third group formed a separate clade, spaced from the isolates of the second group, being the closest to the samples from France, Luxembourg, and United Kingdom. The tree constructed according to the results of phylogenetic analysis of the S gene is shown in Figure 1.

**Figure 1.**
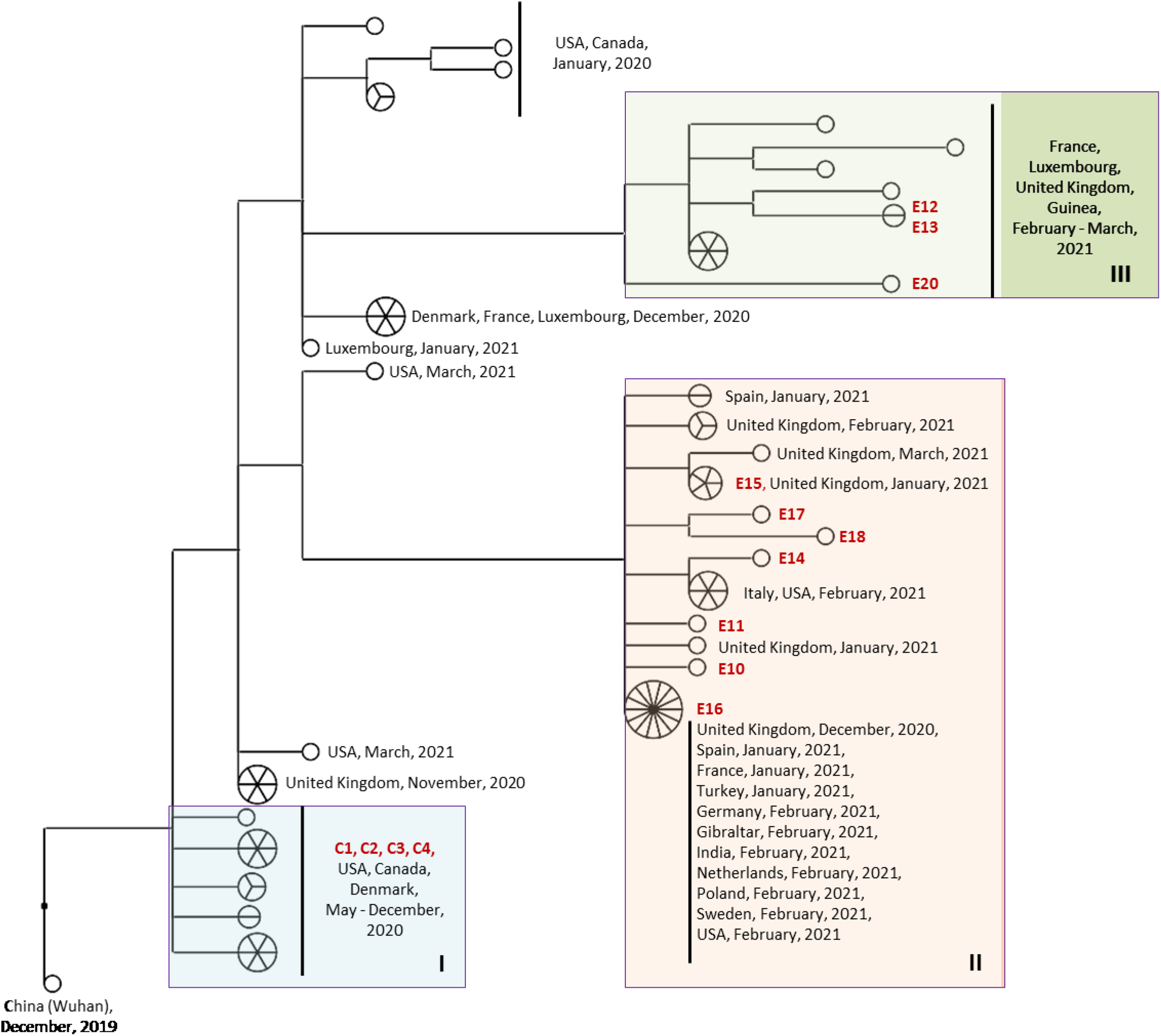
Phylogenetic tree based on the nucleotide sequences of the S gene. The studied samples were divided into three groups (highlighted in color and marked with Roman numerals): I – samples of 2020 not related to known genovariants of epidemic significance; II – samples of 2021, Alpha genotype, or close to it; III – samples from 2021 carrying a common set of mutations in the S gene. The samples examined are highlighted in red.

Since four samples dated 2020, did not have any characteristic features by the nucleotide sequence of the S gene that would allow us to infer their origin, genome-wide sequencing was carried out, followed by the construction of a phylogenetic tree. Mutations in the genome of these strains are shown in Table 3.

**Table 3.**
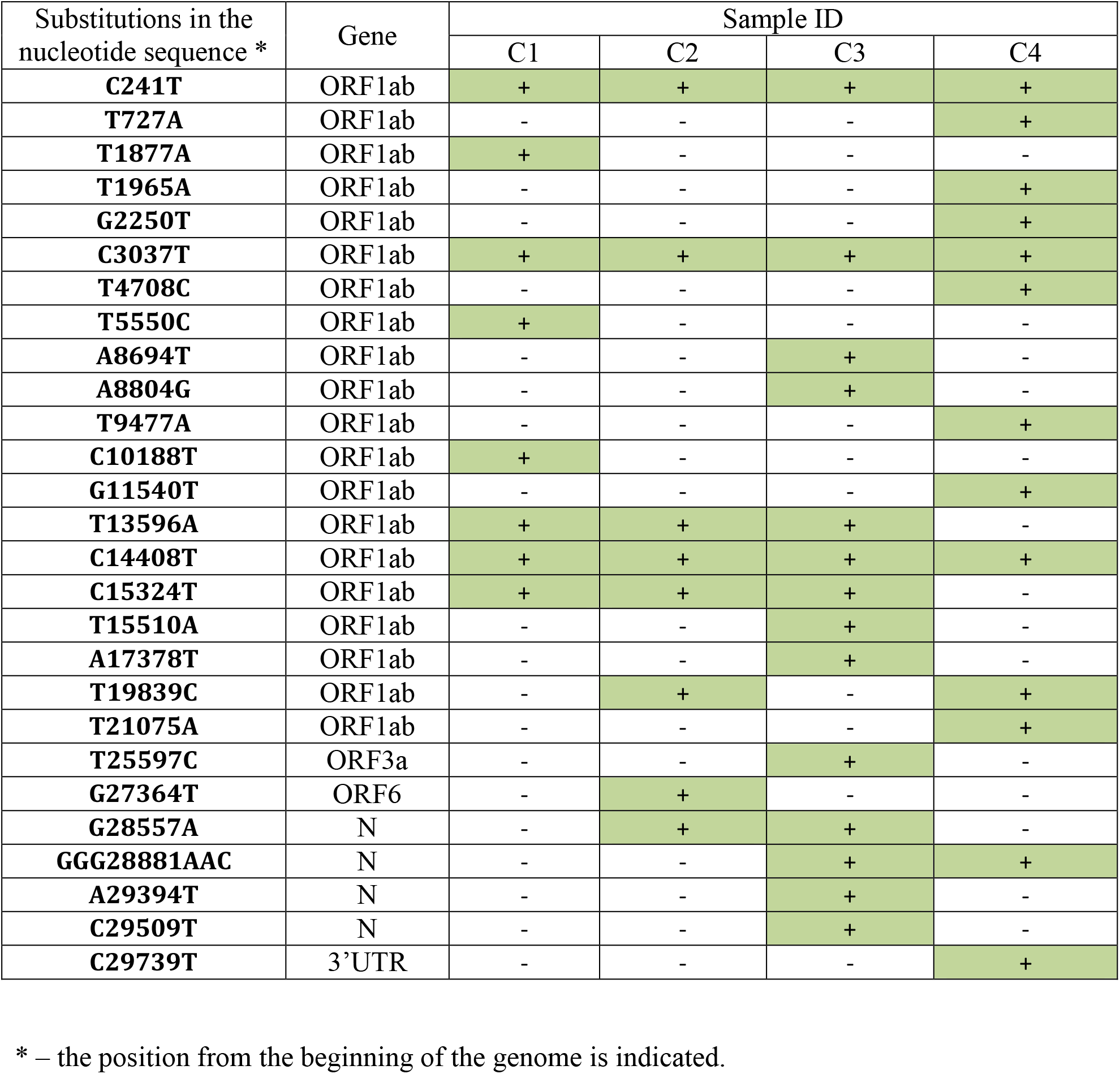
Mutations in the genomes of samples from Guinea. The S gene region is not shown in the table.

The results of sequencing the complete genomes showed that the samples of 2020 belonged to two phylogenetic clades – GR and G. The first clade includes samples C1 and C2 taken in May 2020 in Kolaboui and Sangaredi (Boke prefecture). Both samples belonged to the GR cluster, which does not have a clear geographical confinement. The strains of this cluster were isolated on the territory of the states of Western Europe, North America and Asia. Samples C3 and C4 (June 2020) belonged to clade G and showed high phylogenetic affinity for the strains circulating in Western Europe. So, in phylogenetic construct, these isolates were located in the clade consisting exclusively of genovariants registered in France and / or Switzerland (Figure 2).

**Figure2.**
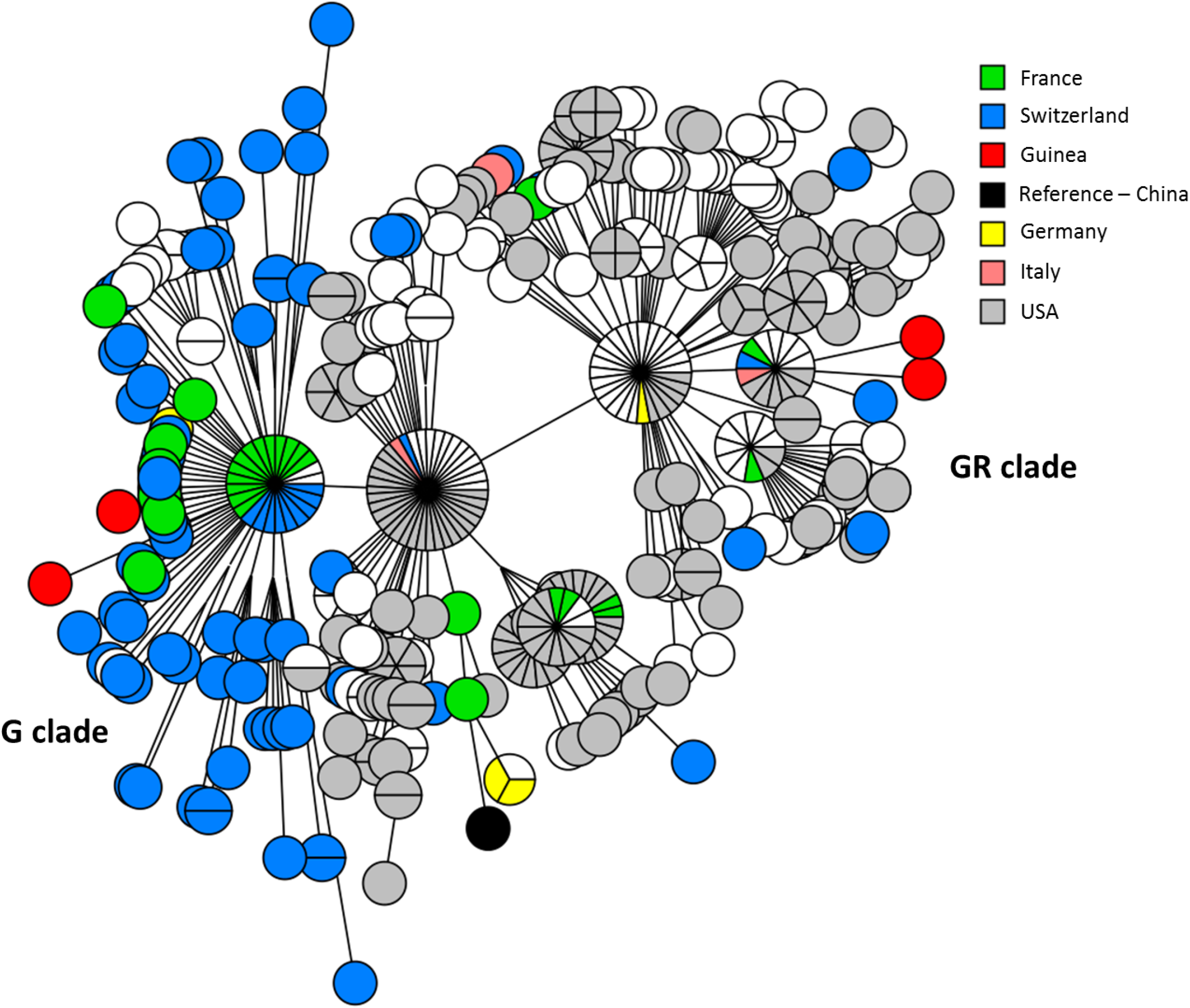
Phylogenetic tree constructed based on sequencing of complete genomes of strains circulating in 2020. Geographical affiliation is shown in color.

Given the high genetic heterogeneity, as well as the different time setting, we can talk about at least four independent imports of coronavirus infection into the territory of the Republic of Guinea. Since the isolates of 2020 demonstrated affinity to two clusters, GR and G, it can be assumed that (before the border closure) at least two independent imports of SARS-CoV-2 genovariants took place in March 2020. And since immediately after the announcement of the COVID-19 outbreak the borders of Guinea became closed, then the internal transmission of infection before the reopening of the borders was precisely through those imported strains.

In confirmation of the epidemiological data, the phylogenetic closeness of the C3 and C4 samples (cluster G) to the samples from France and Switzerland indicates the importation of coronavirus infection from the countries of Western Europe. The strains of the GR cluster (samples C1 and C2) circulating in Guinea at that time were also mainly distributed in Western Europe. Thus, based on the results of our analysis, we have shown that the initial importation of COVID-19 into Guinea took place along at least two independent routes, which are most likely associated with the countries of Western Europe.

After the recommencing of international traffic, despite the anti-epidemic measures being taken, the import of new types of genovariants could not be avoided. So, the Alpha variant already circulated in Guinea at least since the beginning of March 2021 (the date of sampling of E10 is March 02, 2021). Thereat, at the same time, strains phylogenetically distant from the Alpha variant circulated in the same territory (sub-prefecture of Kindia).

Thus, the identification of four SARS-CoV-2 variants, phylogenetically close to the strains circulating outside Guinea, but at the same time phylogenetically distant from each other, suggests at least four independent COVID-19 imports to Guinea – two of which were carried out in March 2020, and two no later than March 2021.

## Conclusions

Based on the above laid out data, the following conclusions can be formulated:

1. The SARS-CoV-2 isolates circulating in the Republic of Guinea are genetically heterogeneous and belong to different phylogenetic groups.
2. Modern genovariants are replacing the originally imported ones.
3. The importation of COVID-19 into the territory of the Republic of Guinea was carried out at least through 4 independent routes.
4. The initial routes of import are connected with the countries of Western Europe.

## Ethical statement

Ethical approval to conduct the study was not needed. Biological specimens were collected as part of the public health emergency response of the Ministry of Health in the Republic of Guinea; therefore, consent for sample collection was waived. Personal information was anonymized.

## References

1. On-line https://www.worldometers.info/coronavirus/

2. A. Rambaut, E. C. Holmes, Á. O’Toole, V. Hill, J. T. McCrone, C. Ruis, L. du Plessis & O. G. Pybus. A dynamic nomenclature proposal for SARS-CoV-2 lineages to assist genomic epidemiology. Nat Microbiol 5, 1403–1407 (2020). https://doi.org/10.1038/s41564-020-0770-5

3. T. Li, T. Huang, C. Guo, A. Wang, X. Shi, X. Mo, et al. Genomic variation, origin tracing, and vaccine development of SARS-CoV-2: A systematic review. Innovation (N Y). 2021 May 28; 2(2):100116. doi: 10.1016/j.xinn.2021.100116

4. W.T. Harvey, A.M. Carabelli, B. Jackson, R.K. Gupta, E.C. Thomson, E.M. Harrison, et al. SARS-CoV-2 variants, spike mutations and immune escape. Nat Rev Microbiol. 2021 Jun 1; 1–16. doi: 10.1038/s41579-021-00573-0

5. S.B. Kadam, G.S. Sukhramani, P. Bishnoi, A. A. Pable, V.T. Barvkar. SARS-CoV-2, the pandemic coronavirus: Molecular and structural insights. J Basic Microbiol. 2021 Mar; 61(3):180–202. doi: 10.1002/jobm.202000537

6. D.J. Benton, A.G. Wrobel, C. Roustan, A. Borg, P. Xu, S.R. Martin et al. The effect of the D614G substitution on the structure of the spike glycoprotein of SARS-CoV-2. Proc Natl Acad Sci U S A. 2021 Mar 2; 118(9):e2022586118. doi: 10.1073/pnas.2022586118

7. P. Majumdar, S. Niyogi. SARS-CoV-2 mutations: the biological trackway towards viral fitness. Epidemiol Infect. 2021 Apr 30; 149:e110. doi: 10.1017/S0950268821001060

8. Qianqian L., Jiajing W., Jianhui N., et al. The Impact of Mutations in SARS-CoV-2 Spike on Viral Infectivity and Antigenicity. Cell, 2020, 182, P. 1284–1294.e9, https://doi.org/10.1016/j.cell.2020.07.012

9. Laffeber C., Koning K., Kanaar R., Lebbink J.H.G. Experimental Evidence for Enhanced Receptor Binding by Rapidly Spreading SARS-CoV-2 Variants. Journal of Molecular Biology. 2021, 433, 167058, https://doi.org/10.1016/j.jmb.2021.167058

10. On-line https://www.who.int/en/activities/tracking-SARS-CoV-2-variants/

11. E. Volz, V. Hill, J. T. McCrone, A. Price, D. Jorgensen, Á. O’Toole et al. Evaluating the Effects of SARS-CoV-2 Spike Mutation D614G on Transmissibility and Pathogenicity. Cell. 2021 Jan 7; 184(1):64–75.e11. doi: 10.1016/j.cell.2020.11.020

12. N.G. Davies, S. Abbott, R.C. Barnard, C.I. Jarvis, A.J. Kucharski, J. D. Munday et al. Estimated transmissibility and impact of SARS-CoV-2 lineage B.1.1.7 in England. Science. 2021 Apr 9; 372(6538):eabg3055. doi: 10.1126/science.abg3055

13. B. Zhou, T.T.N. Thao, D. Hoffmann, A. Taddeo, N. Ebert, F. Labroussaa et al. SARS-CoV-2 spike D614G change enhances replication and transmission. Nature. 2021 Apr; 592(7852):122–127. doi: 10.1038/s41586-021-03361-1

14. S.B. Hudson, V. Kolte, A. Khan, G. Sharma. Dynamic tracking of variant frequencies depicts the evolution of mutation sites amongst SARS-CoV-2 genomes from India. J Med Virol. 2021 Apr; 93(4):2534–2537. doi: 10.1002/jmv.26756

15. M. Shey, J. C. Okeibunor, A. A. Yahaya, B. L. Herring, O. Tomori, S. O. Coulibaly, et al. Genome sequencing and the diagnosis of novel coronavirus (SARS-COV-2) in Africa: how far are we? Pan Afr Med J. 2020 Jun 9; 36:80. doi: 10.11604/pamj.2020.36.80.23723

16. A. Otu, E. Agogo, B. Ebenso. Africa needs more genome sequencing to tackle new variants of SARS-CoV-2. Nat Med. 2021 May; 27(5):744–745. doi: 10.1038/s41591-021-01327-4

17. Russia-Guinea: results and prospects of cooperation in the field of sanitary-epidemiological welfare provision / edited by A. Yu. Popova, Doctor of Medical Sciences, Professor; V.V. Kutyrev, Academician of the Russian Academy of Sciences, Doctor of Medical Sciences, Professor. – Saratov: “Amirit”, 2020. – 272 p.

18. R. Lama, S. Keita, A.A. Kritsky, E.V. Kolomoets, V.K. Konomou, Ya. Yu. Itskov, et al. Anti-epidemic measures in the Republic of Guinea in the fight against COVID-19 in 2020. Proceedings of the International scientific and practical conference on combating new coronavirus infection and other infectious diseases / edited by A. Yu. Popova, Doctor of Medical Sciences, Professor; V.V. Kutyrev, Academician of the Russian Academy of Sciences, Doctor of Medical Sciences, Professor. – Saratov: “Amirit”, 2020. p 130 -132.

19. Plan Nationale De Preparation et De Riposte Contre L’Epidemie de Coronavirus (CoVid-19). Fevrier 2020, Conakry, Guinee.

20. Rapport de Situation. Sitrep N<u>o</u>415. 27/05/2021. ANSS. Guinea, Conakry.

21. A.A. Kritsky, E.V. Kolomoets, K. Sou, S.A. Yakovlev, V.K. Konomou, Ya. Yu. Itskov, et al. Transmission of coronavirus infection in the sub-prefecture of Kindia (Guinea) in the period between march and august 2020. Proceedings of the International scientific and practical conference on combating new coronavirus infection and other infectious diseases / edited by A. Yu. Popova, Doctor of Medical Sciences, Professor; V.V. Kutyrev, Academician of the Russian Academy of Sciences, Doctor of Medical Sciences, Professor. – Saratov: “Amirit”, 2020. p 114–116.

